# Highly efficient chondrogenic differentiation of human iPSCs and purification via a reporter allele generated by CRISPR-Cas9 genome editing

**DOI:** 10.1101/252767

**Authors:** Shaunak S. Adkar, Chia-Lung Wu, Vincent P. Willard, Amanda Dicks, Adarsh Ettyreddy, Nancy Steward, Nidhi Bhutani, Charles A. Gersbach, Farshid Guilak

## Abstract

The differentiation of human induced pluripotent stem cells (hiPSCs) to prescribed cell fates enables the engineering of patient-specific tissue types, such as hyaline cartilage, for applications in regenerative medicine, disease modeling, and drug screening. In many cases, however, these differentiation approaches are poorly controlled and generate heterogeneous cell populations. Here we demonstrate robust cartilaginous matrix production in three unique hiPSC lines using a highly efficient and reproducible differentiation protocol. To purify chondroprogenitors produced by this protocol, we engineered a *COL2A1-GFP* knock-in reporter hiPSC line by CRISPR-Cas9 genome editing. Purified chondroprogenitors demonstrated an improved chondrogenic capacity compared to unselected populations. The ability to enrich for chrondroprogenitors and generate homogenous matrix without contaminating cell types will be essential for regenerative and disease modeling applications.

## Introduction

Chondrocytes produce and maintain the hyaline articular cartilage that lines the surface of diarthrodial joints. Conditions that lead to articular cartilage degeneration, such as arthritis, arise from dysregulated chondrocyte homeostasis, resulting in chondrocyte apoptosis, hypertrophic differentiation of articular chondrocytes, and aberrant activation of inflammatory cascades (Martel-Pelletier et al., 2016; Tonge et al., 2014; van der Kraan and van den Berg, 2012). Ultimately, these changes lead to pain and loss of joint function that necessitates total joint replacement with an artificial prosthesis (Katz, 2006). Many cell-based strategies for cartilage repair and regeneration are designed to preserve the joint (Brittberg et al., 1994; Craft et al., 2015; Estes and Guilak, 2011; K. Lee et al., 2007; Shimomura et al., 2018). Human induced pluripotent stem cells (hiPSCs) are promising cell sources for cartilage regenerative therapies and *in vitro* disease-modeling systems due to their pluripotency and unlimited proliferation capacity (Takahashi et al., 2007; Yamanaka, 2012). Furthermore iPSCs provide a means of developing patient-specific or genetically-engineered cartilage for applications in drug screening for disease-modifying osteoarthritis drugs (DMOADs) (Adkar et al., 2017). Therefore, methods to rapidly and efficiently differentiate hiPSCs into chondrocytes in a reproducible and robust manner will be critical for their application in joint repair.

An important goal of such protocols is to minimize variability in hiPSC differentiation potential, which may arise from characteristics of the donor and/or reprogramming method (Kilpinen et al., 2017; Kim et al., 2011; Nishizawa et al., 2016). Although several approaches have been employed to generate articular chondrocytes from PSCs, most differentiation protocols have been based on trial-and-error delivery of growth factors without immediate consideration of the signaling pathways that direct and inhibit each stage of differentiation. Accordingly, chondrogenic differentiation is often dependent on the specific cell lines used, (Johnstone et al., 2012) and broad application of iPSC chondrogenesis protocols has not been independently demonstrated with multiple cell lines.

Recently, critical insights from developmental biology have elucidated the sequence of inductive and repressive signaling pathways needed for PSC lineage specification to a number of cell fates (Loh et al., 2016; Oldershaw et al., 2010; Umeda et al., 2012). By reproducing these reported signaling pathways *in vitro*, in combination with existing chondrogenic differentiation approaches, we sought to establish a rapid and highly reproducible protocol for hiPSC chondrogenesis that is broadly applicable across various hiPSC lines. Furthermore, as hiPSC differentiation processes are inherently “stochastic” and can often produce heterogeneous cell populations over the course of differentiation, we hypothesized that purification of committed chondroprogenitors would improve hiPSC chondrogenesis. Previously, enrichment of mouse iPSC-derived chondroprogenitors (CPs) resulted from isolating cells expressing *Col2a1*, which encodes for type II collagen – an important structural constituent of articular cartilage (Diekman et al., 2012; Saito et al., 2013; Willard et al., 2014). However, this transgenic approach is not feasible for human iPSCs, which motivated our pursuit of gene editing methods to create a knock-in reporter of collagen II production at the human *COL2A1* locus.

In this study, we demonstrate the development and application of a step-wise differentiation protocol validated in three unique and well-characterized hiPSC lines. We examined gene expression profiles and cartilaginous matrix production during the course of differentiation. To further purify committed CPs, we then used CRISPR-Cas9 genome engineering technology to knock-in a GFP reporter at the collagen type II alpha 1 chain (*COL2A1*) locus to test the hypothesis that purification of CPs could enhance articular cartilage-like matrix production. This type of approach for purification of hiPSC-derived CPs in conjunction with a reproducible differentiation protocol may facilitate the development of *in vitro* chondrogenesis platforms for disease modeling and drug screening.

## Results

### Step-wise differentiation of hiPSCs into chondroprogenitor cells

To establish a standardized protocol for hiPSC chondrogenesis, we optimized growth factor and small molecule concentrations using established principles of PSC differentiation along mesodermal lineages (Loh et al., 2016) as described in **Figure 1A**. To validate our differentiation approach, we measured expression of transcription factors representative of various stages of development with qRT-PCR and monitored cell morphology at multiple time points in three hiPSC lines (BJFF, ATCC, and RVR). (**Figure 1B-G and S1**). Over the course of differentiation, we observed a gradual decrease in expression of the pluripotency markers octamer-binding protein 4 (*OCT4*) and nanog homeobox (*NANOG*) (**Figure 1B**). Stage-specific expression of transcription factors was observed between days 1-3. At day 1, expression of an anterior primitive streak (PS) marker, mix paired-like homeobox (*MIXL1*), spiked in each cell line (**Figure 1C**). Subsequently, a spike in mesogenin 1 (*MSGN1*) expression at day 2 marked paraxial mesoderm specification, and early somitogenesis initiated at day 3, as evinced by upregulation of transcription factor 15 (*PARAXIS*) (**Figure 1D-E**). Interestingly, SRY-box 9 (*SOX9*) expression was highly elevated at both paraxial mesoderm and sclerotome stages as compared to hiPSC stage for ATCC and RVR lines (**Figure 1F**). Furthermore, platelet derived growth factor receptors (*PDGFR*)α and PDGFRβ gradually increased their expression levels over the course of mesodermal differentiation but PDGFRα was down-regulated, whereas PDGFRβ maintained up-regulated, in chondroprogenitor (CP) cells (**Figure 1F**). The CP cells also expressed the highest levels of collagen type II alpha 1 chain (*COL2A1*) but only a moderate increase of aggrecan (*ACAN*) as compared to hiPSCs for all three cell lines examined (**Figure 1G**).

**Figure 1.**
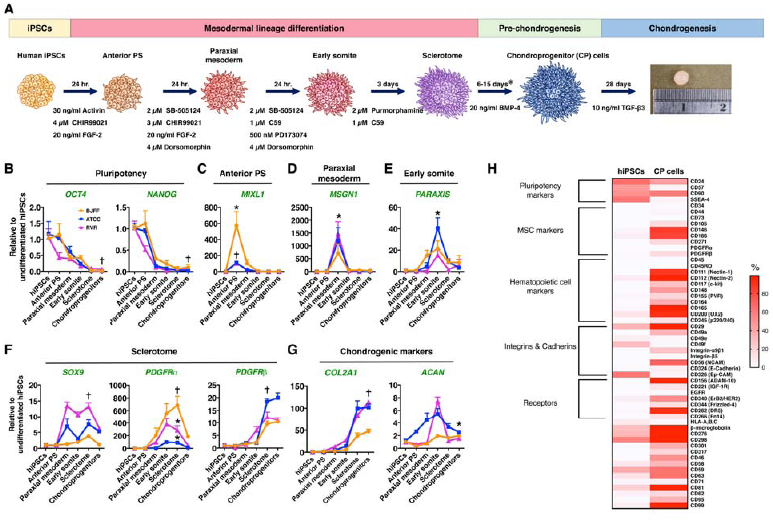
Chondrogenesis of hiPSCs through the Paraxial Mesoderm Lineage. **(A)** Stepwise of differentiation of hiPSCs toward of paraxial mesoderm and chondrocyte-like cells. Gene expression levels of major transcriptional factors for (**B**) pluripotency, (**C**) Anterior PS, (**D**) Paraxial mesoderm, (**E**) Early somite, (**F**) Sclerotome, and (**G**) Chondroprogenitor cells. (**H**) Heat map of representative surface marker expression of hiPSC and Chondroprogenitor (CP) cells of RVR cell line. (**B**-**G**) n = 3-4 experiments per cell stages per cell line (N=3 lines). Mean ± SEM. One-way ANOVA for each cell line using differentiation stage as factor. *p < 0.05 and †p < 0.01 compared to hiPSC stage. For *MSGN1* expression level at paraxial mesoderm stage, only the RVR-iPSC line is significantly different from its hiPSC stage. For *ACAN* expression level at chondroprogenitor stage, only RVR is not significantly different from its hiPSC stage. PS: primitive streak. Data points represent means and error bars signify SEM.

### Characterization of surface markers of hiPSCs and CP cells

Upregulation of chondrogenic markers in CPs suggested that this stage may be appropriate for further chondrogenic differentiation. The RVR-iPSC line was assessed for surface marker expression levels at iPSC and CP cell stages (**Figure 1H**). The hiPSCs and CP cells exhibited distinct expression patterns of surface proteins (**Supplemental Table 1)**. hiPSCs exhibited a surface marker profile characteristic of primed hiPSCs (CD90^+^/CD24^+^/SSEA-4^+^/CD57^+^/CD45^-^). Pluripotency-specific markers SSEA-4/CD57/CD24 decreased in CP cells, and the CP cells displayed a moderate increase in the surface markers CD105, CD146, CD166, and CD271. Interestingly, we also observed that CP cells expressed surface proteins that are often absent on mesenchymal stem cells (MSCs) such as CD56, CD111, CD112, and CD117.

### Chondrogenic gene expression during chondrogenesis

After mesodermal specification and pre-chondrogenesis of hiPSCs, we further differentiated cells in chondrogenic pellet cultures with TGF-β3 supplementation. At days 7, 14, and 28, we assessed chondrogenic gene expression. Chondrogenic markers, particularly SOX9 and *COL2A1*, were significantly up-regulated at early stages of chondrogenesis, while the expression of *ACAN* level was not increased until 14 days post-induction of chondrogenic differentiation (**Figure 2A-C**). These findings suggested that *COL2A1* could serve as critical marker and purification criterion for chondrogenic cells. These three chondrogenic markers remained highly elevated at day 28 of pellet culture as compared to hiPSC stage. We also observed that both hypertrophic chondrocyte marker (collagen type X, *COL10*) and fibroblast matrix gene (collagen type I, *COL1*) were increased at day 28 of chondrogenesis (**Figure 2D-E**).

**Figure 2.**
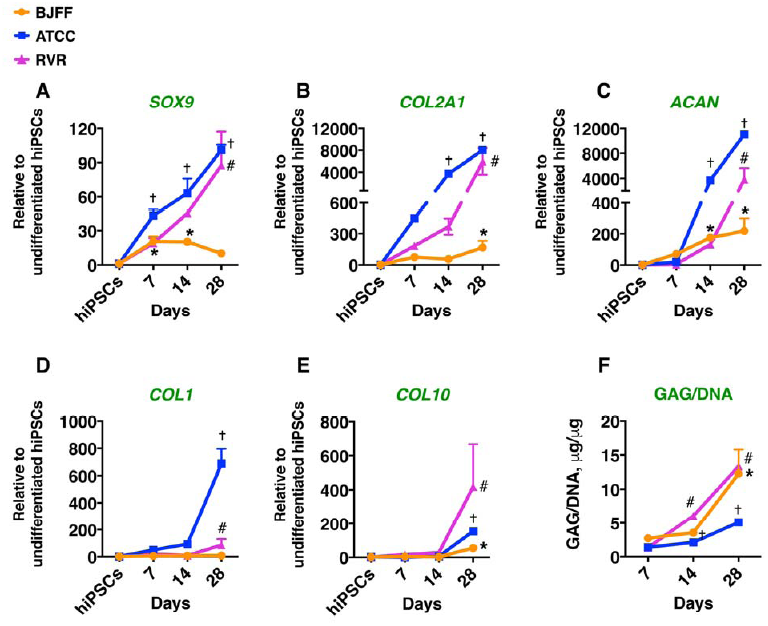
Pellet Culture Time Course. Gene Expression Profiles of **(A**)*SOX9*, (**B)** *COL2A1*, (**C**) *ACAN*, (**D**) *COLIA1*, and (**E**) *COL10A1 and* (**F**) GAG/DNA ratio of the pellets at various time points during 28 day chondrogenic pellet culture at various time points during 28 day chondrogenic pellet culture. n = 3-4 each time point per cell line. Mean ± SEM. One-way ANOVA for each cell line using differentiation stage as factor. *p < 0.05 and †p < 0.01 compared to hiPSC stage.

### Histological and biochemical analyses of chondrogenesis

Safranin-O and hematoxylin staining of day 28 pellets of each cell line showed high amounts of proteoglycan production and the presence of round chondrocyte-like cells within the matrix (**Figure 3**). Additional biochemical analysis of pellets during chondrogenesis demonstrated increasing GAG/DNA ratio over time for each cell line. The ATCC-iPSC line produced less matrix compared to RVR‐ and BJFF-iPSC (**Figure 2F**).

**Figure 3.**
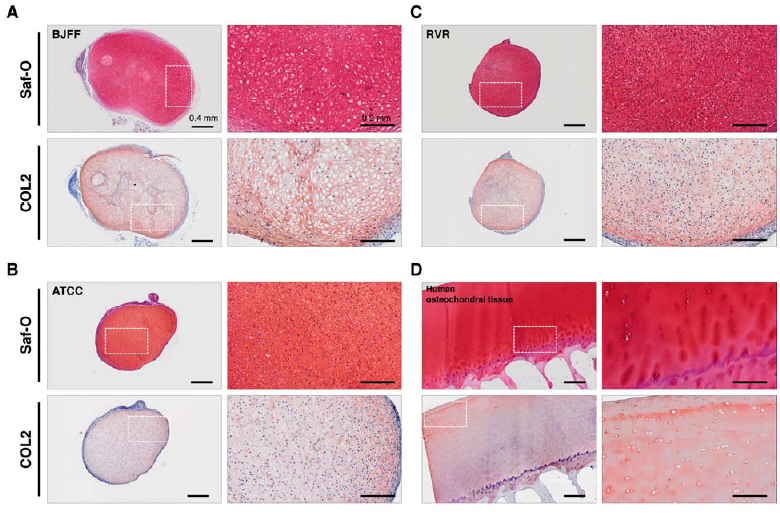
Representative safranin-O (Saf-O) and COL2 staining images of day 28 pellets for **(A**)BJFF (**B)** ATCC, and (**C**) RVR cell line. (**D**) Human osteochondral tissue was used as positive control for both staining. The area of the white dash-square was magnified to reveal chondrocyte-like cell morphology within the pellet for each cell line. Scale bar = 0.4 mm and 0.2 mm for left and right panel, respectively.

### Generation and validation of a COL2A1-GFP knock-in reporter line

While successfully validating a reproducible chondrogenic differentiation protocol applicable to multiple hiPSC lines, we observed some formation of non-cartilaginous tissue from pellet cultures system within each cell line, attesting to heterogeneity inherent in PSC differentiation (**Figure 3A**). Therefore, we sought to further improve cartilaginous matrix production by purifying CP cells prior to chondrogenic differentiation. Using genome editing with CRISPR-Cas9, we integrated an EGFP reporter sequence downstream of the *COL2A1* reading frame, separated by a P2A linker peptide sequence, in RVR hiPSCs (**Figure 4A**). Co-integration of a floxed puromycin resistance cassette allowed enrichment of successfully targeted hiPSCs, and subsequent excision of this cassette left a single loxP scar upstream of the 3’ untranslated region (UTR). Clonal populations were isolated from single cells and successful targeting of a 2A-EGFP sequence at the 3’ end of the *COL2A1* reading frame was confirmed via junction PCR and Sanger sequencing (**Figure S3**).

To functionally validate reporter activity, we activated endogenous *COL2A1* by *SOX9* lentiviral overexpression. Fluorescence imaging revealed clusters of bright round GFP-positive cells after 21 days, comprising roughly 5% of the total population (**Figure 4B** and **4C**). Reporter and unedited RVR lines exhibited comparable activation of *COL2A1* expression, suggesting that the addition of 2A-EGFP and a loxP scar upstream of the 3’ UTR did not significantly alter *SOX9-*dependent *COL2A1* activation (**Figure 4D**). The GFP+ population exhibited significant enrichment (40-fold) of *COL2A1* transcript over unsorted cells; conversely, the GFP-population exhibited decreased levels of *COL2A1* transcripts (**Figure 4D**). Similar trends were observed for expression of other chondrogenic markers such as *ACAN*, *TRPV4*, and *COL9A1* (**Figure 4E-G**).

**Figure 4.**
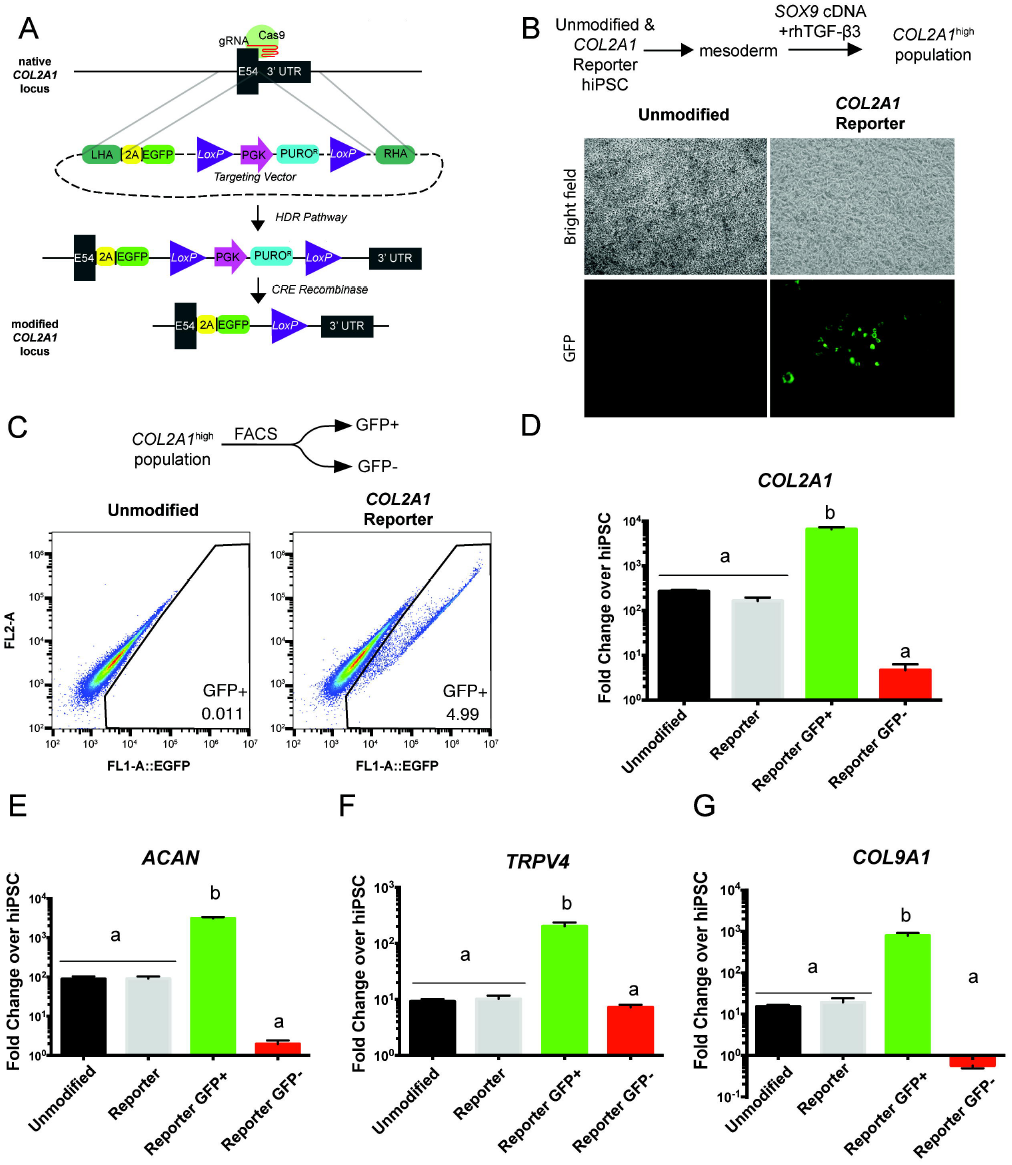
CRISPR-Cas9 mediated knock-in of EGFP Reporter at the *COL2A1* Locus in hiPSCs. (A) Schematic depicting generation of the reporter line. A gRNA overlapping the *COL2A1* stop codon directed Cas9 to generate a double-strand break, which was repaired with a targeting vector containing the 2A-EGFP reporter and a puromycin resistance cassette (PURO^R^) flanked by ∼700 bp homology arms upstream and downstream of the stop codon. After selection of clones with Puromycin, a CRE recombinase expression plasmid was transfected to excise the PURO^R^ cassette, leaving a residual loxP scar. (B) Brightfield and fluorescence imaging of reporter line after *SOX9-*mediated activation of *COL2A1* (C) FACS sorting of GFP+/− populations after *COL2A1* activation in the reporter line and unmodified controls. (D) Fold-change of *COL2A1* expression in unmodified, reporter, GFP+, and GFP-populations relative to expression in hiPSC controls. (E-F) Fold-change of chondrocyte markers *ACAN, TRPV4*, and *COL9A1* in unmodified, reporter, GFP+, and GFP-populations relative to expression in hiPSC controls. Groups not sharing the same letter are statistically different by one-way ANOVA (p<0.05). Error bars represent mean ± SEM (n=3 independent experiments). LHA, RHA: Left Homology Arm, Right Homology Arm

### Application of optimized directed differentiation protocol to the *COL2A1* reporter line

After validating the utility of the reporter in purifying cell populations enriched for *COL2A1* expression, we applied our directed differentiation protocol to the reporter line and observed reporter activation over time (**Figure 5A**). At D0 of differentiation, the COL2A1-GFP reporter line shows no fluorescence when compared to the unmodified line, suggesting the lack of leakiness in reporter activity (**Figure 5Ai**). GFP fluorescence in the reporter line becomes evident as early as D3, during differentiation to paraxial mesoderm (**Figure 5Aii**) and the GFP florescence intensity continues to increase at D6 (**Figure 5Aiii**), finally becoming a distinct GFP+ population by D9 and 12 (**Figure 5Aiv and v**), at the CP cell stage as described previously. At D15 we begin to observe the emergence of a brighter GFP population, which clearly bifurcates into two distinct GFP populations (high and low) at D21 (**Figure 5Avi and vii, Figure 6A**). Mean fluorescence intensity (MFI) of the overall population increased steadily over time (**Figure S4**). To characterize the sensitivity of the reporter in the context of directed differentiation, we sorted various GFP populations: GFP^-^, GFP^+^, GFP^bottom^(bottom 10^th^ percentile of GFP^+^), and GFP^top^(top 10^th^ percentile of GFP^+^), and assayed *COL2A1* transcript levels compared to the unsorted population at multiple time points (**Figure 5B**). Consistent with the intensifying GFP signal over time, we observed a steady increase in *COL2A1* in the unsorted population. GFP^-^ and GFP^bottom^ populations, gated at a constant MFI, exhibited steady levels of *COL2A1* transcript over time (∼100 and 200-fold, respectively). Cell sorting allowed for isolation of cells with significantly greater *COL2A1* transcript levels in the GFP^+^ and GFP^top^ populations compared to unsorted cells (**Figure 5B**).

**Figure 5.**
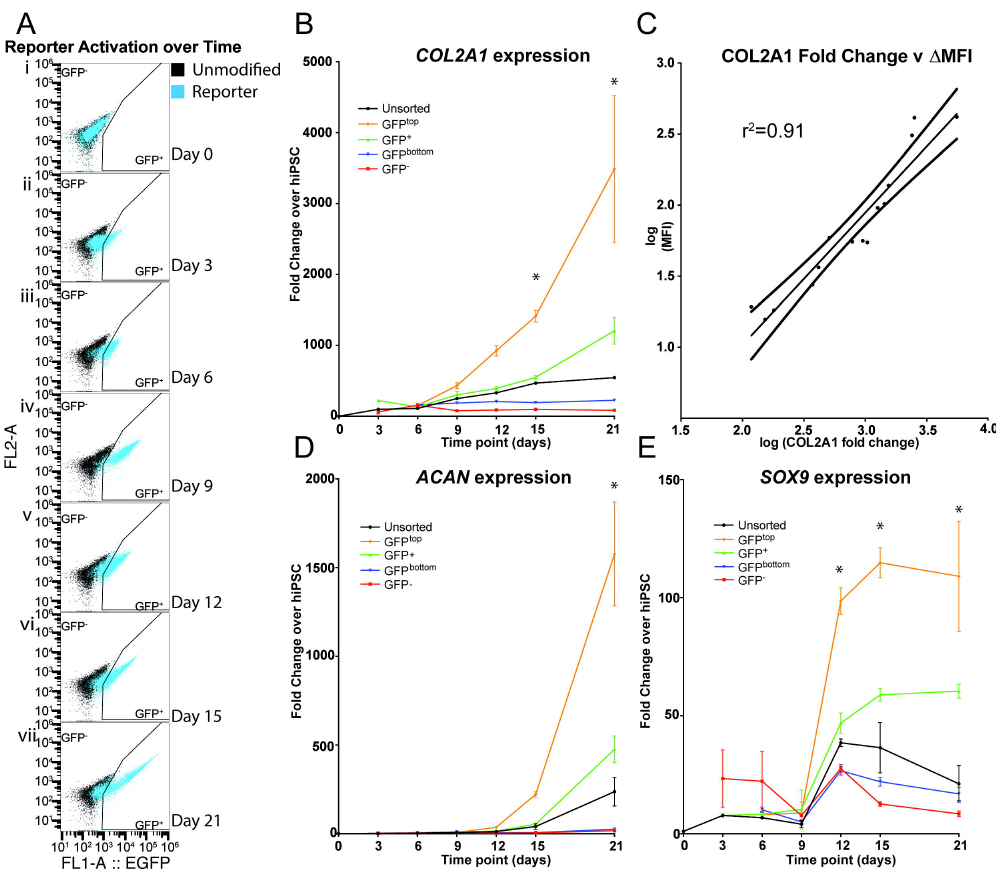
Time course of Directed Differentiation Applied to Reporter Line. (A) FACS sorting of GFP^+/-^ population at several time points. Beginning at D6 the top (GFP^top^) and bottom (GFP^bottom^) 10^th^ percentiles of GFP+ cells were sorted. (B) Differential *COL2A1* transcript enrichment in various GFP populations over the course of differentiation. (C) Scatter plot showing correlation of *COL2A1* transcript enrichment and Mean Fluorescence Intensity (MFI) in GFP^top^ samples. Dashed lines represent 95% confidence intervals. (D,E) Differential *ACAN and SOX9* transcript enrichment in GFP populations over the course of differentiation. Two-way ANOVA with Tukey’s *post hoc* comparison for each chondrogenic marker demonstrated significance along both time and population factors (p<0.05). Asterisks (*) represent significant enrichment (p<0.05) of GFP^high^ compared to each other population at a given time point. Error bars represent mean ± SEM (n=3 independent experiments).

To assay the utility of reporter fluorescence to serve as a surrogate for *COL2A1* expression, we compared the MFI and *COL2A1* fold enrichment of the GFP^top^ population at each measured time point. MFI of GFP^top^ cells was highly correlated to *COL2A1* transcript enrichment (r^2^=0.91) (**Figure 5C**). Taken together, these results suggest that our knock-in reporter is sensitive to relatively small changes in *COL2A1* expression and functions with high fidelity over a wide dynamic range.

GFP^+^ and GFP^top^ populations also exhibited significant enrichment of other chondrocyte-associated transcripts, with robust *ACAN* activation and steady up-regulation of *SOX9* from D12 to D21 (**Figures 5D and 5E**). We also examined expression of Runt-related transcription factor 2 (*RUNX2)*, a marker for hypertrophic or osteogenic differentiation, during the course of differentiation (**Figure S4**). Although we observed enrichment of *RUNX2* in the GFP^top^ at D21, we did not observe activation of its downstream targets including in alkaline phosphatase(ALP), Matrix Metallopeptidase (MMP)-13 and COL10A1 (**Figure S4**). These findings imply that increased *RUNX2* expression by itself may not indicate hypertrophic or osteoblastic differentiation in the GFP^top^ cells. Moreover, transcripts of *COL1A1*, a fibroblastic and osteogenic marker, were significantly depleted in the GFP^+/top^ compared to unsorted controls (**Figure S4**).

### Purification of chondroprogenitors improves cartilaginous matrix production

Given the generation of two distinct GFP populations at D21 (termed GFP^low/high^) of monolayer differentiation (**Figure 6A**) and differential expression of chondrogenic markers within these populations (**Figure 5B, D, and E**), we selected this time point to isolate committed CP cells. GFP^low^ and GFP^high^ populations, along with an intermediate GFP^mid^ population, were sorted and either directly used for chondrogenic pellet culture (P0 pellets) or expanded up to 5 passages prior the induction of chondrogenesis (P1-P5 pellets) (**Figure 6A**). We evaluated pellets after 28 days of culture histologically and biochemically (**Figures 6B-E**). Pellets created from GFP^high^ cells at all passages (P0-P5) were larger than those created from unsorted and GFP^low/mid^ populations, showed homogeneous matrix rich in glycosaminoglycan (GAG) and were void of other tissue types (**Figure 6B and S5A**). Biochemical analysis confirmed the histology with significantly higher GAG/DNA ratio in the GFP^high^ pellets compared to unsorted ones. Interestingly, chondrogenesis of GFP^mid^ cells improved with expansion, suggesting that this population may have been a less differentiated population compared to GFP^high^. Meanwhile GFP^low^ populations failed to produce cartilaginous matrix from P0-P5 (**Figure 6C**).

**Figure 6.**
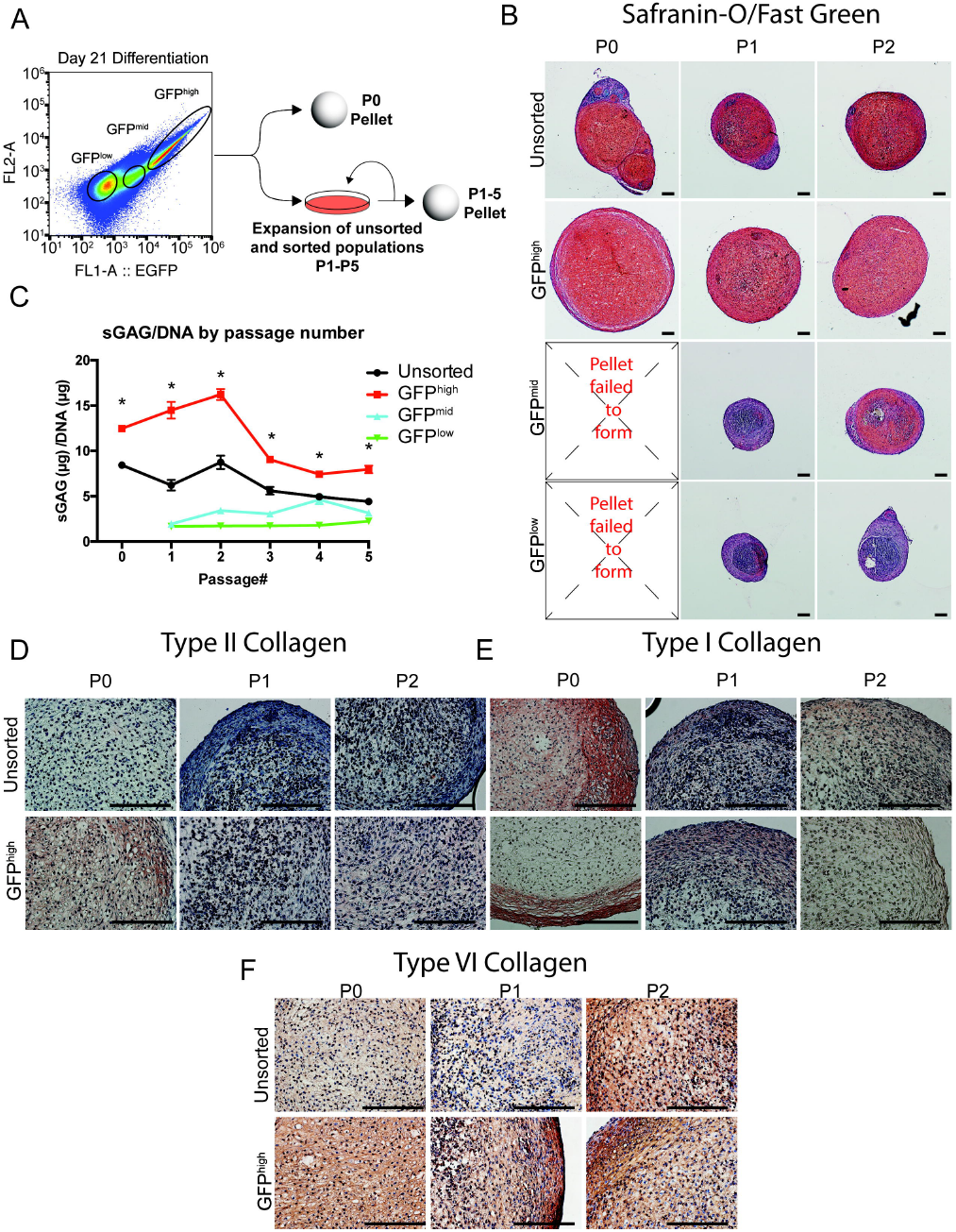
Evaluation of Chondrogenic Differentiation Capacity of Purified Chondroprogenitors. (A) Schematic of Experimental Design. D21 GFP^high/low/mid^ populations were isolated and cultured as aggregates or seeded for expansion. Aggregates were generated from expanded cells at each passage up to 5 passages. (B) Safranin-O/Fast Green stain of aggregates from unsorted and GFP populations after FACS (P0) and 1 or 2 passages (P1 and P2, respectively) Scale bar (200 μm) (C) Biochemical analysis of GAG content normalized to DNA content in unsorted and GFP populations. Two-way ANOVA with Tukey’s *post hoc* comparisons for each chondrogenic marker demonstrated significance along both time and population factor (p<0.05). Asterisks (*) represent significant enrichment (p<0.05) of GFP^high^ compared to each other population at a given passage. Error bars represent mean ± SEM (n=3 independent experiments). (D-F) Type II Collagen (D), Type I Collagen (E), and Type VI Collagen (F) immunohistochemistry (IHC) of pellets derived from unsorted cells and GFP^high^ cells at P0-P5. Scale Bar (200 μm)

While the presence of type I collagen predominates in fibrocartilage, hyaline articular cartilage is characterized by the presence of type II collagen and the lack of type I collagen. Immunohistochemistry analysis of collagen composition demonstrated enrichment of type II collagen in P0-P2 pellets generated from GFP^high^ cells compared to the unsorted cells (**Figure 6D**). P0 pellets from unsorted cells stained diffusely for type I collagen, and this was limited to the periphery of the tissue from GFP^high^ P0 pellets (**Figure 6E**). Type VI collagen, another component of articular cartilage, was also enriched in P0-P2 pellets derived from GFP^high^ cells (**Figure 6F**). Together, this suggests that purification of GFP^high^ cells specifies the generation of matrix with qualities similar to hyaline cartilage rather than fibrocartilage. Pellets derived from passaged GFP^high^ CPs (P1-P5) had less type II collagen accumulation as compared to P0 pellets (**Figure 6D, Figure S5D**). In addition, late passage GFP^high^ pellets (P3-P5) exhibited increased type I collagen accumulation compared to unsorted controls (**Figure S5E**).

## Discussion

Articular chondrocytes develop *in vivo* from a group of mesenchymal cells derived from mesoderm and ectoderm (Decker et al., 2014; Marianne E Bronner, 2012; Oldershaw et al., 2010; Yamashita et al., 2015b). While craniofacial cartilage is generated from the ectoderm-derived neural crest, articular cartilage of the appendicular and axial skeleton is derived from paraxial and lateral plate mesoderm, respectively. Recent advances in developmental biology and the establishment of hPSC differentiation “roadmaps” can be applied to the *in vitro* generation of various musculoskeletal tissues through the specification of different mesodermal lineages (Craft et al., 2015; Loh et al., 2016; Oldershaw et al., 2010). While it is known that chondrocytes can be derived from paraxial mesoderm during embryonic development *in vivo*, the reproducible differentiation of hiPSCs to chondroprogenitors (CPs) has remained a major challenge to cartilage tissue engineering and arthritis disease modeling. Here, we describe a highly efficient, stepwise protocol for *in vitro* chondrogenesis through the paraxial mesodermal lineage that is consistent among multiple hiPSC lines and further validated using a knock-in collagen II reporter system.

Our approach successfully recapitulates mesodermal differentiation, as evidenced by the sequential stage-specific up-regulation of mesodermal master transcription factors including *MIXL1*, *MSGN1* and *PARAXIS*. Interestingly, the findings of down-regulation of *PDGFRα* as well as *SOX9*, but not *PDGFRβ*, in chondroprogenitor (CP) cells in response to BMP-4 treatment suggests possible interactions among PDGFRα, SOX9, and BMP signaling pathways during CP cell commitment. Indeed, PDGFRαexpression level has been reported to be involved in regulating BMP-Smad1/5/8 signaling and induction of osteogenesis in MSCs (Li et al., 2014). Furthermore, using conditional deletion animal models, Lim at al recently demonstrated that BMP-Smad4 signaling controls pre-cartilaginous mesenchymal condensation in a SOX9 independent manner (Lim et al., 2015).

CP cells expressed high levels of CD146 and CD166 but moderate to low levels of CD271, CD73, CD44, and stage specific embryonic antigen (SSEA)-4. These results suggest that the CP cells derived via our differentiation protocol are distinct from MSCs, which generally express these markers (CD271, CD73, CD44, SSEA4) (Lv et al., 2014). Moreover, our finding of low CD44 levels in the CP cells is consistent with the results of a recent study showing that CD44 is expressed at a low level in pre-chondrocytes as compared to mature chondrocytes (Wu et al., 2013). However, the results of surface marker analyses should be interpreted with caution, as surface markers do not necessarily reflect cell function and as stem and/or progenitor cells can be highly plastic, and lineage-origin dependent.

The cartilaginous matrix produced by the pellets showed consistent staining for type II collagen and GAG, but was negative for type I collagen, similar to hyaline articular cartilage. Indeed, diffuse expression of type VI collagen further suggests the production of immature “neocartilage”. Prolongation of chondrogenesis in pellet culture may improve maturation of hiPSC derived-chondrocytes. Our protocol generates articular cartilage by extended treatment with TGF-β3, which has been shown to serve as an important step in the generation of articular cartilage from adult stem cells as well as human PSCs (Craft et al., 2015; Diekman et al., 2012; Johnstone et al., 1998; Yamashita et al., 2015). Extended treatment with BMP-4 in pellet culture on the other hand may promote a more hypertrophic phenotype (Craft et al., 2015).

Our study describes the generation of articular cartilage-like tissue derived through the paraxial mesodermal lineage, but the subtle differences between human chondrocytes derived from paraxial/lateral plate mesoderm or ectoderm remain unclear. Future investigation of hiPSC chondrogenesis along alternative developmental lineages, coupled with comprehensive genomic, transcriptomic, and/or proteomic analysis may facilitate identification of functional differences at multiple stages of differentiation. This information may be important for functional investigation of genetic arthritis susceptibility variants, which may cause joint-specific pathology (Pan et al., 2014; Zhang et al., 2015).

With the widespread availability of iPSCs and gene editing technologies, we now have the ability to assay the functional effects of disease susceptibility variants on developmental and pathological phenotypes (Ding et al., 2013; Gupta et al., 2017; Musunuru, 2013; Shi et al., 2017; Smith et al., 2014; Takahashi et al., 2007; Warren et al., 2017). Genome-wide association studies (GWAS) and candidate association studies have identified many genetic variants that may be involved in pathologies of articular cartilage, such as rheumatoid arthritis and osteoarthritis (arcOGEN Consortium et al., 2012; Zhu et al., 2016). Follow-up analyses of these genetic variants has been challenging in iPSC-derived CPs due to the difficulty of isolating this cell population. Indeed, the heterogeneity of *in vitro* chondrogenesis may preclude our understanding of the mechanisms by which genetic variation can impact cartilage development and arthritis progression. Previous attempts to purify iPSC-derived CPs rely on many rounds of sorting multiple surface markers, using reporter constructs that are activated by minimal promoter regions of chondrocyte markers, or physical dissection of self-aggregating chondroprogenitors (Diekman et al., 2012; Umeda et al., 2015; Wu et al., 2013; Yamashita et al., 2013; 2015a). During chondrocyte differentiation, *COL2A1* expression is regulated by concerted coordination of promoter regions, distal and intronic enhancers, and UTRs (Han and Lefebvre, 2008; Jash et al., 2012; Lefebvre et al., 1997). Therefore, the knock-in reporter approach described in this study enables precise monitoring of endogenous *COL2A1* expression during differentiation and purification of a homogeneous CP population.

High-throughput disease modeling and drug screening platforms of cartilage injury and repair will benefit from the ability to produce large quantities of homogeneous cartilaginous aggregates from purified CPs (Adkar et al., 2017; J. Lee et al., 2015; Umeda et al., 2015; Willard et al., 2014; Yamashita et al., 2015b). Our laboratory and others have demonstrated the utility of CPs as an expandable depot of cells for subsequent chondrogenesis (Diekman et al., 2012; Umeda et al., 2015). We show that the day 21 purified CPs (GFP^high^ population) maintain improved chondrogenic capacity over the course of 5 passages compared to controls. In this manner, small molecule and cytokine screens can be readily applied to tissue derived from purified CPs in a high throughput manner (Willard et al., 2014). In addition to drug screening, CRISPR screening technology has recently become a powerful tool to identify coding and non-coding genetic elements that perturb reporter or surface marker expression or affect viability (Joung et al., 2017; Klann et al., 2017; Parnas et al., 2015). Since our reporter enables purification of CP cells, it can also facilitate such screens to identify novel targets necessary or inhibitory for chondrogenesis. Validation of identified hits may provide targets to further improve hiPSC chondrogenesis protocols.

The future of novel arthritis therapeutics lies in the application of hiPSC chondrogenesis to regenerative medicine and arthritis disease modeling (Adkar et al., 2017; Craft et al., 2015). The standardization of chondrogenesis protocols that follow developmental pathways will allow comparisons to be made between studies and improve reproducibility. By applying signaling logic aimed at activating and repressing specific developmental pathways, we can reliably guide hPSC differentiations along mesodermal lineage bifurcations (Loh et al., 2016; 2014). This work shows that establishment of a standardized chondrogenesis protocol combined with purification of CP cells can produce a homogenous population of chondrocyte-like cells with improved matrix quality. Future studies investigating the effects of genetic or environmental perturbations on hiPSC-derived cartilage-like tissue will benefit from the current differentiation method.

## Experimental Procedures

### hiPSC Lines and Cell Culture

Three human iPSC lines were used in the current study: BJFF.6, ATCC^®^ACS-1019 and RVR-iPSC (J. Lee et al., 2015; 2012). Both BJFF.6 (BJFF) and ATCC^®^ACS-1019 (ATCC) were derived from foreskin fibroblasts of male newborn infants using sendai virus. BJFF was obtained from iPSC core at Washington University in St. Louis, and the cells were maintained on vitronectin (VTN-N; Fisher Scientific, #A31804) coated plates in Essential 8 Flex medium (E8; Life Technologies, A2858501). The ATCC line was purchased from ATCC (ATCC^®^ ACS1019^TM^), and the cells were maintained on CellMatrix Basement Membrane Gel (ATCC, ACS-3035) in Pluripotent Stem Cell SFM XF/FF medium (ATCC, ACS-3002). RVR-iPSCs were retrovirally reprogrammed from BJ fibroblasts and characterized previously (J. Lee et al., 2012) The cells were maintained on vitronectin in mTeSR1 medium (Stemcell Technologies, #05857). Media were changed daily and cells were passaged at 80-90% confluency.

### Mesodermal Lineage Differentiation

Mesodermal differentiation medium: induction of hiPSCs into mesodermal lineages is based on a protocol reported by Loh et al with slight modifications (Loh et al., 2016). In brief, when hiPSCs reached 70% confluency, the culture medium was switched to mesodermal differentiation medium composed of IMDM GlutaMAX^TM^ (IMDM, Fisher Scientific, #31980097) and Ham’s F12 Nutrient Mix (F12, Fisher Scientific, #11765054) with 1% chemically defined lipid concentrate (Gibco, #11905031), 1% insulin / human transferrin / selenous acid (ITS+; Corning, #354352), 1% penicillin / streptomyocin (P/S; Gibco, #15140122), 450 μM 1-thioglycerol (Sigma, #M6145), and 1 mg/ml polyvinyl alcohol (Sigma, P8136-250G).

Induction of anterior primitive streak (PS) (day 0): hiPSCs were rinsed with wash medium consisting of IMDM, F12 and 1% PS, and then fed with mesodermal differentiation medium supplemented with 30 ng/ml of Activin A (Stemgent, #03-0001), 4 μM of CHIR99021 (CHIR; Stemgent, #04-0004), and 20 ng/ml human fibroblast growth factor (FGF; R&D Systems, 146 aa recombinant protein, #233FB001MG/CF) for 24 hrs.

Induction of paraxial mesoderm (day 1): after a brief wash with DMEM/F12, cells were fed with mesodermal differentiation medium supplemented with 2 μM SB-505124 (SB5; Tocris, #3263), 3 μM CHIR, 20 ng/ml FGF, 4 μM dorsomorphin (DM; Stemgent, #04-0024) for 24 hrs.

Induction of early somite (day 2): after a brief wash with DMEM/F12, cells were fed with mesodermal differentiation medium supplemented with 2 μM SB5, 4 μM DM, 1 μM Wnt-C59 (C59; Cellagen Technology, #C7641-2s), and 500 nM PD173074 (PD17; Tocris, #3044) for 24hrs.

Induction of sclerotome (day 3 – day 5): after a brief wash with DMEM/F12, cells were fed with mesodermal differentiation medium supplemented with 2 μM purmorphamine (Pur; Stemgent, #04-0009) and 1μM C59 for 3 days. Medium was changed daily.

Induction of chondroprogenitor cells (day 6 to day 12-21): after a brief wash with DMEM/F12, cells were fed with mesodermal differentiation medium supplemented with 20 ng/ml human bone morphogenetic protein 4 (BMP-4; R&D Systems, #314-BP-01M). Media was changed daily.

### Chondrogenic Differentiation

Monolayer cultured CP cells were dissociated with ethylenediaminetetraacetic acid (EDTA) at 37°C, and EDTA was diluted with an equal amount of medium composed of DMEM/F-12, GlutaMAX^TM^ (DMEM/F12; Gibco, #10565-018), 10% fetal bovine serum (FBS; Atlanta Biologicals, #S11550), 1% ITS+, 55 μM 2-Mercaptoethanol (2-ME, Gibco, #21985), 1% MEM non-essential amino acids (NEAA; Gibco, #11140050), and 1% P/S. The cells were then collected and centrifuged for 5 minutes at 300 x g. After aspirating the supernatant, the cells was re-suspended in chondrogenic medium composed of DMEM/F-12, 1% FBS, 1% ITS+, 55 ìM â-mertcaptoethanol, 100 nM dexamethasone (DEX; Sigma-Aldrich, #D4902), 1% NEAA, 1% P/S, 10 ng/ml human transforming growth factor beta 3 (TGF-β3; R&D Systems, 24-3B3), 50 μg/ml L-ascorbic acid 2-phosphate (ascorbate; Sigma-Aldrich, #A8960), and 40 μg/ml L-Proline (proline; Sigma-Aldrich, #P5607), and 300,000 − 500,000 CP cells were then pelleted for chondrogenesis for 28 days. Chondrogenic medium was changed every three days.

### Histology

Pellets were fixed in 10% formalin for overnight and stored in 70% ethanol at 4°C. They were then embedded in paraffin wax and sectioned at 8 μm thickness. Sections were stained with Safranin-O/hematoxylin standard protocol to reveal proteoglycan matrix.

### Immunohistochemistry

To perform immunohistochemistry of type II collagen (II-II6B3, 1:1), type VI collagen (70RCR009×, Fitzgerald, 1:200), or type I collagen (90395, abcam, 1:800), sections were treated with xylenes and a decreasing ethanol series, incubated for 5 min with Digest-All 3 pepsin digestion (Invitrogen), treated with 30% peroxidase, blocked with goat serum, incubated with primary antibody for 1 h at room temperature, incubated with secondary antibody (ab97021 for type II and type I collagen, ab6720 for type VI collagen, Abcam, 1:500) for 30 minutes, treated with HRP streptavidin and AEC Red Single (Histostain-Plus BS, Invitrogen), counterstained with hematoxylin, and mounted with Clearmount (Invitrogen). Pig and human osteochondral sections were used as both positive and negative controls.

### Biochemical Analysis

Pellets were rinsed with DPBS, digested overnight at 65°C in 200 μl of papain solution consisting of 125 μg/ml papain, 100 mM sodium phosphate, 5 mM EDTA, and 5 mM L-cysteine hydrochloride at 6.5 pH, then stored at −20°C. Double-stranded DNA content was quantified using the PicoGreen^TM^ assay (Invitrogen, #P7589). Total sulfated glycosaminoglycan (s-GAG) content was measured using the dimethymethlyene blue (DMMB) assay using bovine chondroitin sulfate (Sigma Aldrich, #C9819) as a standard.

### Gene Expression

Cells in monolayer and pellets were rinsed with DPBS. Monolayer cells were lysed in 600 μl of Buffer RL (Norgen Biotek) and pellets were snap-frozen in 350 μl of Buffer RL which were then stored at −80°C. Pellets were homogenized using a miniature bead beater. The RNA was isolated using the Total RNA Purification Kit according to the manufacture’s recommendations (Norgen Biotek, #17200). Reverse transcription was performed using SuperScript^TM^ VILO^TM^ Master Mix (Thermo Fisher Scientific, #11755500) per the manufacturer’s instructions. Quantitative RT-PCR was performed on the QuantStudio 3 (Biosystems) and CFX96 Real Time System (Biorad) using Fast SYBR^TM^ Green Master Mix (Thermo Fisher Scientific, #4385616) according to the manufacturer’s protocol. Fold changes were calculated using the ΔΔC_T_ method relative to hiPSCs as the reference time point and TATA-box-binding protein (TBP) as the reference gene. Gene expression was probed using the primer pairs listed in the **Supplemental Table 2.**

### Flow Cytometric Analysis for Surface Markers

Surface markers of hiPSCs and CP cells of RVR-iPSCs were characterized by 361 PE-conjugated monoclonal antibodies (LEGENDScreen Human PE Kit, #700007) according to manufacturer’s instructions. Stained cells were fixed and analyzed by LSRII (BD Biosciences).

### Fluorescence Activated Cell Sorting

At various time points during differentiation, cells were dissociated as described above. Pelleted cells were re-suspended in sort buffer containing PBS, 10% FBS, 2x P/S/F, and 1 U/mL DNase I (Thermo, EN0525)). Cells were sorted on a Sony SH800S Cell Sorter (Sony Biotechnology) using a 130 μm sorting chip.

### Genome Editing and Clonal Isolation

To generate the *COL2A1* reporter line, RVR-iPSCs were dissociated with accutase and 3x10^6^ cells were re-suspended in 200 μL complete mTeSR. 6 μg of gRNA/Cas9-2a-GFP expression vector and 3 μg of *COL2A1* targeting vector were added (see **Supplemental Experimental Procedures**). Cell suspension was placed into a 4 mm cuvette and transfected using the Gene Pulser XL (BioRad) electroporation system. Transfected cells were plated in a matrigel-coated 10cm dish in complete mTeSR supplemented with 10μM Rock Inhibitor (Y-27632). Selection began the following the day with 0.5 μg/mL Puromycin. After roughly 10 days puromycin-resistant colonies were transfected with a CRE recombinase expression vector to remove the floxed selection cassette. Transfected cells were expanded and plated at low density for clonal isolation(180 cells/cm^2^). After 10-14 weeks resulting clones were mechanically picked and expanded for PCR screening of targeting vector integration using the following primer pair: PuroF2: 5’-GCAACCTCCCCTTCTACGAG; RightOutReverse: 5’-GCAAGGGACACCCAAGAGTA-3’

### Statistical Analysis

To compare matrix production and gene expression levels within each cell line (Figures 1,2, and 4), one-way ANOVA using differentiation stage as the factor with Tukey’s *post hoc* testing was performed. For comparisons between multiple groups at various passages or time points (Figure 5 and 6), a two-way ANOVA using time point/passage and GFP population as factors with Tukey’s *post hoc* testing were performed as appropriate. All statistical analyses were carried out using the Graphpad Prism software. Significance was reported at the 95% confidence level.

## Author Contributions

Conceptualization, S.S.A., C.L.W., V.P.W., C.A.G., N.B., and F.G.; Methodology, S.S.A., C.L.W., V.P.W., C.A.G., and F.G.; Investigation, S.S.A., C.L.W, V.P.W., A.D., A.E., N.S.; Formal Analysis, S.S.A, C.L.W, and V.P.W.; Writing – Original Draft, S.S.A., C.L.W, and V.P.W.; Writing – Review and Editing, S.S.A., C.L.W, V.P.W., A.D., N.B., C.A.G., and F.G.; Supervision, C.A.G. and F.G.; Funding Acquisition, C.A.G. and F.G.

## Acknowledgments

We would like to acknowledge the support by the Nancy Taylor Foundation for Chronic Diseases, the Arthritis Foundation, NIH grants AR061042, AR50245, AR48852, AG15768, AR48182, DK108742, OD008586, GM119914, HG007900, DA036865, AR069085, NSF CAREER Award CBET-1151035, and the Collaborative Research Center of the AO Foundation, Davos, Switzerland.

---

**Supplemental Figure 1, related to Figure 1.** Cell morphology of each stage during mesodermal lineage differentiation for each cell line. Fibroblastic-like cells started to emerge at stage of early somite. Scale bar = 500 μm

**Supplemental Figure 2, related to Figure 2.** Representative Negative control of COL2 stain for (**A**) BJFF cell line and (**B**) human osteochondral tissue. Scale bar = 0.4 mm

**Figure S3. Confirmation of Reporter Integration, related to Figure 4** (A) Junction PCR using primers shown in red to validate site-specific knock-in of EGFP reporter. One primer binds GFP while the other binds 3’ of right homology arm. See Supplemental Experimental Procedure for primer sequences.

**Figure S4, related to Figure 5.** (A) Histograms showing reporter activation over time and production of two distinct GFP populations by Day 21 in independent three replicates of differentiation. (B) Average MFI at each time point. (C) Evaluation of *RUNX2* transcript enrichment in various GFP populations at multiple time points. (D-G) Fold change at Day 21 of hypertrophic markers *COL1A1* (D), *COL10A1* (E), *ALP* (F), *MMP-13* (G) relative to expression in hiPSCs. Error bars represent mean ± SEM (n=3 independent experiments).

**Figure S5, related to Figure 6.** (A) Safranin-O/Fast Green stain of aggregates of unsorted and GFP populations from P3-P5. (B,C) Type II (B) and Type I (C) collagen IHC of pellets derived from P0-P2 GFP^low/mid^ cells. (D,E) Type II (D) and Type I (E) collagen of pellets derived from unsorted and GFP^low/mid/high^ at P3-P5.

